# Self-assembly of nanofilaments in cyanobacteria for protein co-localization

**DOI:** 10.1101/2023.02.12.528169

**Authors:** Julie A. Z. Zedler, Alexandra M. Schirmacher, David A. Russo, Lorna Hodgson, Emil Gundersen, Annemarie Matthes, Stefanie Frank, Paul Verkade, Poul Erik Jensen

## Abstract

Cyanobacteria offer great potential as alternative biotechnological hosts due to their photoautotrophic capacities. However, in comparison to established heterotrophic hosts, several key aspects, such as product titers, are still lagging behind. Nanobiotechnology is an emerging field with great potential to improve existing hosts but, so far, it has barely been explored in microbial photosynthetic systems. Here, we report the establishment of large proteinaceous nanofilaments in the unicellular model cyanobacterium *Synechocystis* sp. PCC 6803 and the fast-growing cyanobacterial strain *Synechococcus elongatus* UTEX 2973. Transmission electron microscopy and electron tomography demonstrated that overexpression of a modified bacterial microcompartment shell protein, PduA*, led to the generation of bundles of longitudinally aligned nanofilaments in *S. elongatus* UTEX 2973 and shorter filamentous structures in *Synechocystis* sp. PCC 6803. Comparative proteomics showed that PduA* was at least 50 times more abundant than the second most abundant protein in the cell and that nanofilament assembly only had a minor impact on cellular metabolism. Finally, we targeted the fluorescent reporter mCitrine to the nanofilaments using an encapsulation peptide that natively interacts with PduA. To our knowledge, this is the first study to apply bacterial microcompartment based nanotechnology in cyanobacteria. The establishment of nanofilaments in cyanobacterial cells is an important step towards cellular organization of heterologous pathways and the establishment of cyanobacteria as next generation hosts.

## Introduction

Nanobiotechnology is an emerging field at the interface of nano- and biotechnology which deals with the synthesis, manipulation and biofunctionalization of structures at the nanometer scale. Conceptually, the ambition to program units at the local scale to achieve desired properties at the global scale intertwines with another fast-developing field, synthetic biology. When applied in a bottom-up approach, synthetic biology concerns itself with building new-to-nature structures and functions with naturally occurring biological units. One area where both fields converge is in their desire to control and exploit nanoscale spatial organization.

The ability to control the spatial organization of biological units has many applications, however one particularly promising use is the scaffolding of biosynthetic pathways. In their native environment, sequential enzymes of a biosynthetic pathway often use structural elements of the cell (e.g. lipid membranes, proteinaceous compartments) to form dynamic supramolecular complexes termed metabolons^1^. The assembly of enzymes into a metabolon can increase substrate channeling, reduce crosstalk with native pathways and accumulation of toxic intermediates, increasing the catalytic efficiency of the pathway. However, when biosynthetic pathways are expressed in a heterologous chassis, the native structural and regulatory elements needed to form the metabolon are often lost, leading to low product titers. To address this, there have been efforts to engineer artificial nanostructures, e.g. scaffolds, within heterologous chassis where enzymes can be co-localized to form metabolic hotspots^2,3^. The vast majority of enzyme scaffolds have used DNA or protein subunits^4^. Examples include the dockerin and cohesin domains deriving from cellulosomes of cellulolytic bacteria^5^, the SH3, PDZ and GBD domains from metazoan cells^6^, affibodies^7^ and DNA origami tiles^8^. While scaffolding technology has been successfully applied in heterotrophic chassis^9^, such as *E. coli* and yeast, it has yet to be developed for photoautotrophic microbes.

Cyanobacteria are a diverse group of photosynthetic bacteria that hold great promise as future production chassis due to their fast growth and ability to fix atmospheric carbon dioxide. However, production titers often lag behind heterotrophic production hosts, in part due to difficulties in efficient enzyme targeting and co-localization inside a densely packed cell structure with two membrane systems. In this study we address this challenge by introducing, for the first time, a synthetic scaffold in cyanobacteria.

Using a modified bacterial microcompartment (BMC) shell protein, we show that nanofilaments can be formed in both coccoid and rod-shaped cyanobacteria and that a heterologous reporter protein can be targeted to these nanofilaments. Finally, growth experiments and a global proteome analysis suggest that nanofilament formation has little impact on native metabolism. The introduction of soluble proteinaceous nanofilaments for spatial organization of heterologous proteins marks a major milestone on the path to making cyanobacteria competitive biotechnological hosts in a future green bioeconomy.

## Methods

### Generation of plasmids

All cloning was performed using *Escherichia coli* NEB 5-alpha. An overview of the design of individual constructs used in this study is shown in Figure 5A and details are given in Supplementary Table 1. The gene sequence of *pduA\** (a derivative of *pduA* from *Citrobacter freundii* with a C-terminal extension)^10^ was inserted into the self-replicative shuttle plasmid pDF-trc^14^ using restriction digest cloning at the *EcoRI* and *HindIII* sites and the generated plasmid was termed pEG001. For co-expression of PduA* and mCitrine, four additional constructs were made by Gibson assembly^39^. The plasmids pAS001 and pAS002 contain synthetic operons of *pduA\** and *mCitrine* with (pAS001) and without (pAS002) a DNA sequence encoding an encapsulation peptide (P18, AA sequence: MNTSELETLIRNILSEQL)^31^ fused with a flexible GGGS linker to the N-terminus of mCitrine. The plasmids pAS003 and pAS004 are controls for expression of mCitrine (pAS003) or P18-mCitrine (pAS004) in the absence of PduA*. All primers used for plasmid generation are specified in Supplementary Table 2. Fragments for Gibson assembly were generated by PCR using Q5 Hot Start High-Fidelity polymerase (New England Biolabs, USA). PCR products were analyzed by agarose gel electrophoresis and bands of the expected size were excised, purified and used for Gibson assembly using HiFi DNA assembly Master Mix (New England Biolabs, USA). Correct assembly of all constructs was confirmed by Sanger sequencing (Eurofins Genomics, Germany).

### Cyanobacterial strains and growth conditions

Two different cyanobacterial species were used in this study: *Synechococcus elongatus* UTEX 2973 and *Synechocystis* sp. PCC 6803 (a glucose-tolerant strain “Kaplan” originally obtained from Prof. Patrik Jones, Imperial College London). All cultures were maintained at 30°C with approximately 50 μmol photons m^-2^ s^-1^ on 1.5% agar plates with BG-11 medium^40^ supplemented with 10 mM 2-tris(hydroxymethyl)-methyl-2-amino 1-ethanesulfonic acid (TES), pH 8.0. For all strains containing expression plasmids, the growth medium was supplemented with 50 μg mL^-1^ spectinomycin. Liquid cultures were grown at 30°C in P4 medium^41^ supplemented with 50 μg mL^-1^ spectinomycin. For electron microscopy and proteome analysis, cells were grown in a high-density cultivation setup (HDC 6.10B starter kit from CellDEG GmbH, Germany). A culture volume of 10 mL was used in 25 mL vessels with CO_2_ supplementation through a membrane at approximately 5% from a 3 M KHCO_3_ and 3 M K2CO_3_ buffer (ratio 4:1). Cultures were illuminated with approximately 50 μmol photons m^-2^ s^-1^ (PCC 6803) or 125 μmol photons m^-2^ s^-1^ (UTEX 2973) continuous illumination. All cultures were induced with 1 mM isopropyl β-D-1-thiogalactopyranoside (IPTG) after 24 h and grown for a total of three days for electron microscopy or four days for proteomics.

### Generation of cyanobacterial strains

Plasmids were introduced into UTEX 2973 and PCC 6803 by conjugation using either biparental (pEG001, pDF-trc) or triparental (pAS001-pAS004) mating. Biparental mating was performed as described previously^42^. For triparental mating, *E. coli* NEB 5-alpha carrying the target plasmid and helper strain *E. coli* HB101 with plasmids pRL433 and pRL623^43^ were grown in LB medium at 37°C and harvested during exponential growth (Optical Density (OD) at 600 nm ≈ 0.5). Cells of both *E. coli* strains were mixed in an equal ratio and incubated at 30°C for 30 min before mixing with cyanobacteria (OD_750 nm_ = 1 of cultures in exponential growth). The mixture was added to a cellulose acetate/cellulose nitrate filter piece on a BG-11 + 5% (v/v) LB agar plate and further treated as for biparental mating^42^. Colonies were analyzed by colony PCR for the presence of the target plasmid using the same method as described in Russo et al. 2019^42^ with the extension time at 72 °C extended to 1.5 min.

### Growth analysis and mCitrine fluorescence monitoring

Cultures were grown in P4-TES CPH medium^42^ supplemented with 50 μg mL^-1^ spectinomycin at 30°C with air bubbling supplemented with 3% CO_2_. Samples were taken twice daily to measure the OD at 750 nm and mCitrine fluorescence (Agilent BioTek Synergy H4 Hybrid Reader, ex. 490 nm, em. 531 nm). UTEX 2973 was inoculated to an OD_750 nm_ of 0.2 in 20 mL P4-TES CPH medium, induced with 1 mM IPTG and incubated for 96 h with 720 μmol photons m^-2^ s^-1^ illumination. PCC 6803 was inoculated to an OD_750 nm_ of 0.4 in P4-TES CPH medium and illuminated with 75 μmol photons m^-2^ s^-1^. After 24 h of growth, cultures were induced with 1 mM IPTG and growth of the cultures was monitored for a total of 96 h.

### Sucrose density gradients

Cultures were grown in the same conditions as for growth analysis. 48 h after induction, an OD_750 nm_ equivalent of 20 (UTEX 2973) or 40 (PCC 6803) was harvested. Samples were prepared for fractionation on a sucrose gradient based on BMC enrichment protocols^44,45^. Cell pellets were resuspended in 2 mL TE lysis buffer (10 mM Tris-HCl pH 7.5, 1 mM ethylenediaminetetraacetic acid (EDTA), 5% Glycerol, 1% protease inhibitor Mix B (SERVA Electrophoresis GmbH, Germany)) and lysed with a Bullet Blender Storm 24 (Next Advance, USA) with 0.15 mm zircodium oxide beads. Subsequently, 1 % n-dodecyl β-D-maltoside (DDM) in TE buffer was added and the mixture incubated for 1 h at 4°C. The samples were spun down at 10 000 × g for 10 min at 4°C to remove cell debris and the supernatant was carefully loaded onto a layered sucrose column, consisting of 60%, 40%, 20%, 15% and 10% sucrose. The sucrose gradients were centrifuged at 200 000 × g for 20 h at 4°C (Sorvall Discovery 90 SE Ultracentrifuge, rotor SW 40). Individual sucrose fractions were then carefully collected using glass Pasteur pipettes and further processed for SDS-PAGE.

### Preparation of cleared lysates and samples for SDS-PAGE

For protein expression analysis, cleared cell lysates were prepared as previously described^42^. Protein content of all samples analyzed by SDS-PAGE was estimated using a Pierce bicinchoninic acid (BCA) protein assay (Thermo Scientific, Germany) using the 96-well plate protocol as recommended by the manufacturer.

### SDS-PAGE, Coomassie or Oriole staining and immunoblotting

SDS-PAGE was performed as previously described^46^. Gels were then either stained with Oriole Fluorescent Gel Stain (Bio-Rad), Coomassie Brilliant Blue R-250 or used for immunoblotting. For immunoblotting, proteins were transferred onto a 0.2 μm nitrocellulose membrane (Protran BA 83, Amersham) by semi-dry blotting and further processed as described previously^46^. For the detection of PduA*, a customized antibody was generated by Agrisera (Sweden). This antibody was raised against the synthetic peptide “CPHTDVEKILPKGI” in rabbit. For the detection of mCitrine by western blotting, a monoclonal Anti-eGFP antibody raised in mouse (MA1.952, Invitrogen) was used.

### Proteomics

Cleared cellular lysates were generated using a modified lysis buffer recipe (20 mM Tris-HCl pH 7.5, 5% glycerol, 0.02% Rapigest SF (Waters)) using the same protocol as used for expression analysis. 2 μL of cleared cellular lysates at a concentration of 5.5 to 8.5 mg mL^-1^ total protein were reduced with 10 mM dithiothreitol and alkylated with 55 mM iodoacetamide. Sequential digestion with Lys-C (Promega) and trypsin (proteomic grade from porcine pancreas, Sigma-Aldrich) was performed to manufacturers’ recommendations followed by desalting using Oasis HLB 1cc 10 mg extraction cartridges. Samples were analyzed using a Dionex Ultimate 3000 series UHPLC system coupled with a hybrid TIMS QTOF (Bruker timsTOF Pro). Peptides were separated prior to MS on an Acclaim PepMap column 100, 75 μm x 50 cm nanoViper C18 3 μm 100A column. Mobile phases A and B were 0.1% formic acid and 80% acetonitrile with 0.08% formic acid, respectively. Peptides were separated at 0.4 mL min^-1^ using an isocratic gradient for 5 min at 5% B, followed by a linear gradient from 5% to 40% B over 210 min, 40% to 80% B over 5 min and held for an additional 5 min before re-equilibration to 5% B over 15 min. The UHPLC was coupled via a CaptiveSpray nano-electrospray source to the mass spectrometer and the capillary heated to 180°C. Data acquisition on the timsTOF Pro used the parallel accumulation serial fragmentation (PASEF) mode. The instrument was operated in positive ion mode with a mass range of 100-17 000 *m/z.* PASEF settings included 10 MS/MS scans at 1.19 s total cycle time, target intensity of 20 000, charge range of 0-5 and active exclusion release after 0.4 min. The spectra were searched against the reference proteome for *S. elongatus* PCC 7942 (UniProt Proteome ID UP000630882) or PCC 6803 (UniProt Proteome ID UP000001425) using Mascot Daemon. The search was performed with a 15 ppm error for peptide masses, whereas 0.5 Da were allowed for the fragmented products. Carbamidomethylation was set as fixed modification and oxidation as variable. The cleavage needed to be at one side of the peptide tryptic. Two missed cleavages were allowed in total. Proteins were considered identified when two unique peptides per protein were matched. Protein abundance was estimated by the exponentially modified protein abundance index (emPAI)^47^. Data processing, statistical analysis (log2 fold change and Student’s t-test) and data visualization was done with the support of Microsoft Excel and R^48^ (“tidyverse”^49^, “hablar”^50^, “EnhancedVolcano”^51^). The mass spectrometry data have been deposited at ProteomeXchange via the XXX partner with dataset identifier XXX.

### TEM and electron tomography

Cells were grown as previously described and harvested by centrifugation at 3000 × g for 5 min. 1 μL of cell pellets were transferred onto copper carrier grids and subjected to high-pressure freezing using a Leica EMPACT2 high-pressure freezer + Rapid Transfer System (RTS) (Leica Microsystems)^52^. The frozen samples were transferred to 1% Osmium tetroxide and 0.1% uranylacetate in acetone and freeze substituted using a Leica EM AFS2 freeze substitution processor at – 90°C with a slow warm up of 5°C per hour up to 0°C (Leica Microsystems). As osmium becomes increasingly reactive at higher temperatures we removed the samples from the AFS2 at 0°C to preserve the “crispiness” of the staining. Samples were then rinsed in pure acetone and infiltrated with increasing concentrations of Epon resin (25%, 50%, 75%, 100%, in acetone), embedded in moulds and heat polymerized. Ultrathin sections of 70 nm were cut using a diamond knife and a UC6 ultramicrotome (Leica Microsystems) and imaged on a Tecnai T12 transmission electron microscope (FEI / Thermo Fisher). For electron tomography, 300 nm sections of the same samples were cut and labelled with fiducials (15 nm gold particle solution) on both surfaces. Tilt series were recorded from −60 to +60° imaged on a Tecnai T20 transmission electron microscope (FEI / Thermo Fisher) and tomograms were reconstructed using IMOD software^53^.

## Results and Discussion

### Expression of a modified Pdu BMC shell protein leads to nanofilament formation in cyanobacteria

PduA* is a derivate of PduA, a shell protein of the propanediol utilization (Pdu) BMC in *Citrobacter freundii^10,11^*. In comparison to PduA, PduA* has a C-terminal extension of 23 amino acids which leads to higher solubility^10^. PduA* is a homo-hexamer, which was found to self-assemble, through non-covalent interactions, into bundles of nanotubes in *Escherichia coli*^10–12^ and *Citrobacter glutamicum^13^.* In order to achieve co-localization of heterologous proteins within the cyanobacterial cell, we first set out to generate a *de novo*, protein-based scaffold structure in the cytoplasm of the cyanobacterial cell utilizing the PduA* protein. This protein was overexpressed under the control of a trc promoter in two different cyanobacterial strains, *Synechococcus elongatus* UTEX 2973 (hereafter UTEX 2973) and *Synechocystis* sp. PCC 6803 (hereafter PCC 6803) using a high expression, self-replicative, plasmid system^14^. PCC 6803 is a widely used, classic cyanobacterial model organism where most metabolic engineering studies have been conducted^15–18^ and UTEX 2973 is a fast-growing strain^19^ with high biotechnological relevance. Further, their contrasting cellular morphologies, coccoid (PCC 6803) versus rod-shaped (UTEX 2973), might give insights into how scaffold assembly is influenced by cellular architecture in cyanobacteria.

After confirming successful transformation by colony PCR (Supplementary Figure 1), analysis of whole cell lysates by SDS-PAGE and Coomassie staining showed a strong additional band around 12 kDa in the cyanobacterial transformants carrying the *pduA\** gene (Figure 1A, pEG001 plasmid). This fits with the predicted molecular weight of PduA* of 11.9 kDa. An antibody, specifically raised for the detection of PduA*, further confirmed that this highly abundant protein was PduA* (Figure 1A).

**Figure 1.**
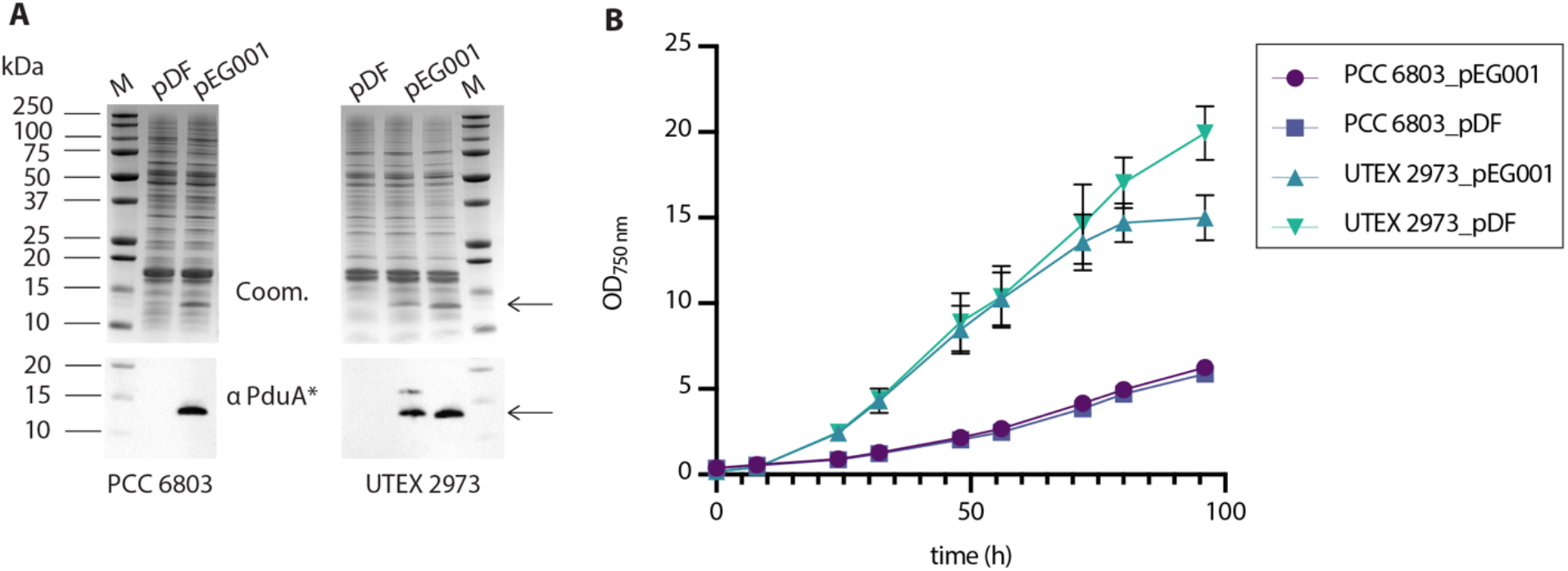
PduA* expression in cyanobacteria. Two different cyanobacteria, *Synechocystis* sp. PCC 6803 and *S. elongatus* UTEX 2973, were transformed with the self-replicating plasmid pEG001 for PduA* expression or an empty control plasmid (pDF). **A** Coomassie stain (Coom.) and anti-PduA* immunoblot (αPduA*) of cleared whole cell lysates. M denotes a molecular weight marker**. B** Growth of generated strains measured by optical density at 750 nm. UTEX 2973 cultures were grown with 720 μmol photons m^-2^ s^-1^ continuous illumination, PCC 6803 strains with 75 μmol photons m^-2^ s^-1^. Data points shown are averages, error bars represent the standard deviation (n = 3).

We then analyzed the growth of both PCC 6803 and UTEX 2973 expressing PduA* (pEG001 plasmid) in comparison to an empty vector control (pDF plasmid) over four days (Figure 1B). In both cyanobacteria, PduA* expression did not significantly impact growth under the experimental conditions tested. However, the PduA* expressing UTEX 2973 strain (UTEX 2973_pEG001) entered stationary phase earlier than the control strain resulting in a lower final biomass yield (Figure 1B). Virtually no differences in growth were observed between the PduA*-expressing PCC 6803 strain (PCC 6803_pEG001) and the empty vector control (PCC 6803_pDF) for the duration of the growth period. The differences in growth observed between PCC 6803 and UTEX 2973 strains were expected given the strain characteristics^20^. Surprisingly, the high level of PduA* expression did not significantly change the growth profile of the recombinant strains. In *Escherichia coli,* PduA* expression was reported to interfere with septation^10,12^ but growth data were not reported, therefore a comparison with our findings was not possible.

Having confirmed that PduA* can be successfully expressed in cyanobacteria, we used transmission electron microscopy (TEM) and electron tomography to investigate whether PduA* expression in cyanobacteria leads to the assembly of cytoplasmic supramolecular structures. Cells were cultured in standard growth conditions, fixed, stained, embedded, and sectioned for TEM imaging.

In UTEX 2973_pDF, a typical ultrastructure of a rod-shaped cyanobacterial cell, with thylakoid membranes aligned concentrically along the plasma membrane, was found in both longitudinal and transversally cut sections of cells (Figure 2D). This structure and size is in line with what was originally described for this strain^19^ and what is known from the closely related model cyanobacterium *Synechococcus elongatus* PCC 7942^21^. In the UTEX 2973_pEG001 strain, additional filamentous structures, distinct from thylakoid membranes, were observed in longitudinal sections (Figure 2A, 2B, Supplementary Video 1, 2). These filamentous structures were similar to what was observed for PduA* expression in *E. coli*^10–12^ and *C. glutamicum^13^* as well as for other hexameric BMC shell proteins overexpressed in *E. coli^22,23^.* In addition, we frequently observed native carboxysomes linearly aligned in central cytoplasmic regions in close proximity to PduA* nanofilaments (Supplementary Video 1, 2, 3, Figure 2A, 2B). This carboxysome alignment is in accordance with reports of β-carboxysome positioning in other rod-shaped cyanobacteria such as the closely related strain *S. elongatus* PCC 7942^24–26^. Transverse sections of UTEX 2973_pEG001 showed bundles of nanotube-like structures (Figure 2C, Supplementary Video 3). In comparison to previous reports from *E. coli*, these nanotube-like structures appear less regularly packed resembling filamentous structures of a mutant version of PduA (V51A)^11^. In addition, highly ordered honeycomb structures were also observed in the UTEX 2973_pEG001 transverse sections (Figure 2C). These structures resemble those formed by PduA^11^ and PduA*^10^ in *E. coli* and may derive from bundles of tightly packed nanofilaments. However, despite appearing distinct from carboxysomes observed in the control UTEX2973_pDF (Supplementary Figure 2), we cannot exclude that these structures are native carboxysomes.

**Figure 2.**
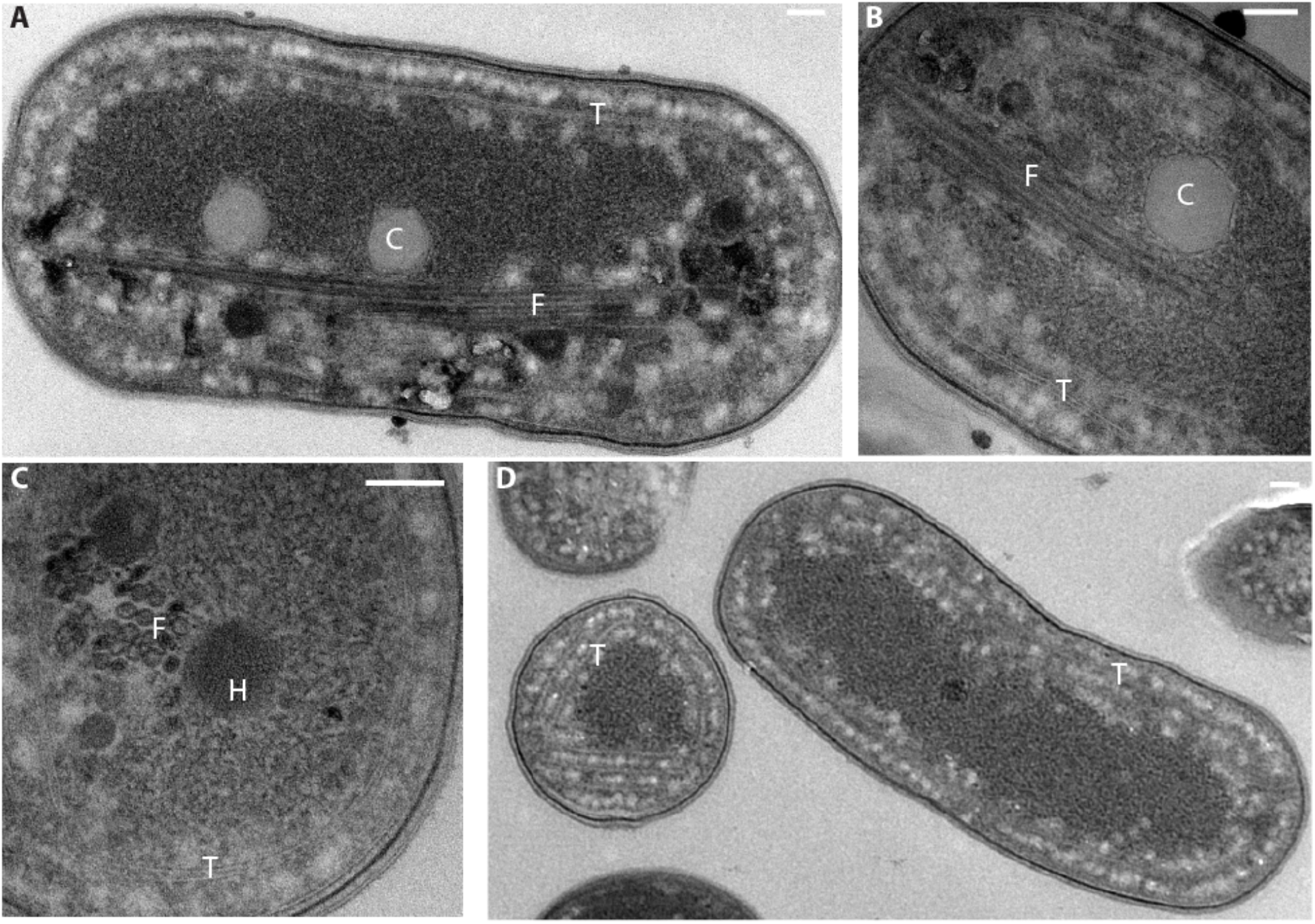
Ultrastructure of PduA* nanofilaments in *S. elongatus* UTEX 2973. Transmission electron micrographs of UTEX2973_pEG001 strain (**A, B, C)** and an empty vector control strain (UTEX 2973_pDF) (**D).** Panel C shows a transverse view of a cell. T denotes thylakoid membranes, C carboxysome, F PduA* filaments and H honeycomb structures. Scale bars: 100 nm.

Overall, based on our TEM micrographs and electron tomography data, we can conclude that PduA* hexamers in UTEX 2973_pEG001 assemble into nanofilamentlike supramolecular structures that span the entirety of the cell. We speculate that the differences found between hexameric BMC nanofilaments in heterotrophic hosts and those formed in cyanobacteria may be due to an interplay with the carboxysomes and elements of the cytoskeleton^27,28^. Further work will be needed to understand these interactions in more detail and characterize the underlying molecular mechanisms.

In PCC 6803_pEG001, the architecture of PduA*-derived supramolecular structures was less clear to determine by TEM than for UTEX 2973_pEG001. Nonetheless, filamentous structures, absent from the empty vector control and distinct from thylakoid membranes, were observed in the PduA* expressing strain (Figure 3A). It is likely that the coccoid cellular architecture of PCC 6803, together with its crowded cellular environment, results in the formation of shorter nanofilaments than what was observed in the rod-shaped UTEX 2973. In some sections, filamentous structures that resemble highly stacked thylakoid membranes, similar to grana stacks in chloroplasts of higher plants, were observed (Supplementary Figure 3). This architecture has not been reported for PCC 6803 thylakoid membranes and might be caused by interference of PduA* with the native cellular architecture, however further evidence is needed for clarification. Overall, the evidence based on TEM imaging clearly demonstrates the formation of nanofilaments. However, it is not sufficient to determine whether the observed filamentous structures in PCC6803_pEG001 form nanosheets or whether these sheets are rolled up into nanotubes as previously postulated for higher order assembly of PduA*^11^.

**Figure 3.**
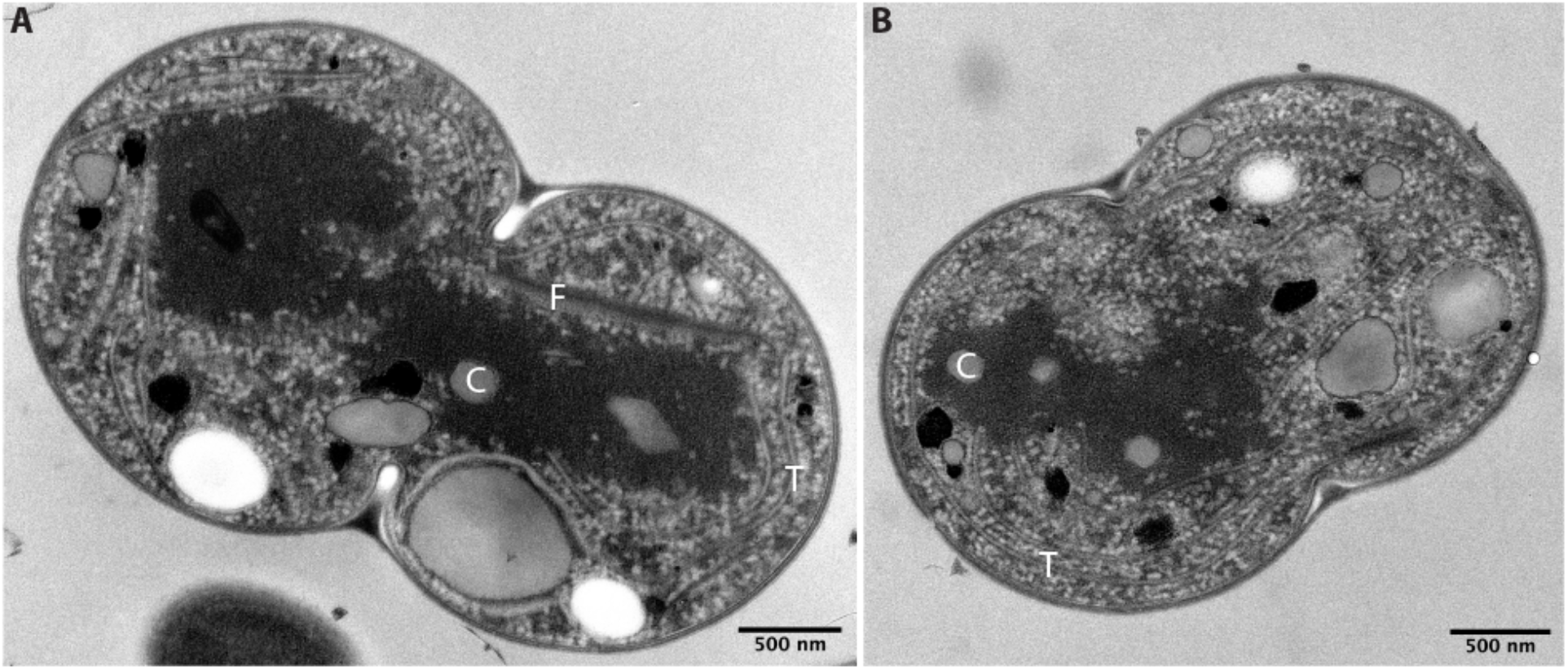
Ultrastructure of PduA* nanofilaments in *Synechocystis* sp. PCC 6803. Transmission electron micrographs of PCC6803_pEG001 (**A)** and an empty vector control strain (PCC6803_pDF) (**B)**. T denotes thylakoid membranes, C carboxysome and F PduA* filaments. Scale bars: 500 nm.

PduA* expression was previously reported to interfere with septation in E. coli^10,12^. We saw evidence for this in UTEX293_pEG001 (Supplementary Figure 4), but not for PCC 6803_pEG001. The differences in PduA* ultrastructure observed between both cyanobacterial strains might be due to an interrelationship of filament assembly and cellular architecture. This may also explain the differences observed in our growth data where no significant impact was found for PCC 6803_pEG001 while UTEX 2973_pEG001 showed a lower final biomass yield (Figure 1). A hypothesis is that the cell-spanning nanofilaments interfere with septation and, consequentially, impact growth. In longitudinal sections of UTEX 2973_pEG001 cells, we also observed transverse cuts of nanofilaments (Supplementary Figure 4). This suggests that filament elongation in UTEX 2973 seems to occur not only along the central longitudinal axis of the cell but in various directions. The underlying factors that determine in what directions PduA* nanofilaments extend in cyanobacteria are yet to be determined.

### Nanofilament formation does not have a major metabolic impact on the cell

Having established that PduA* nanofilaments can be assembled in both cyanobacteria, we proceeded to assess their metabolic impact using a comparative, label-free, shotgun proteomics approach. The most abundant protein identified in both nanofilament-forming strains was PduA*. In both strains the PduA* monomer was identified with 100% protein coverage and was found to be 53 and 63 times more abundant than the second most abundant protein in UTEX 2973 and PCC 6803, respectively (Supplementary Data D1). To evaluate the metabolic impact of nanofilament formation, we proceeded with a comparative analysis. Only proteins identified by at least two tryptic peptides and present in both the nanofilament strains (UTEX 2973_pEG001 and PCC6803_pEG001) and their respective empty vector controls (UTEX 2973_pDF and PCC6803_pDF) were considered. In total, 1008 and 1260 proteins were identified in UTEX 2973 and PCC 6803, respectively (Supplementary Data D1). Differentially expressed proteins were selected from all identified proteins with a cut-off criterium of fold change = 2 and p-value < 0.05 (Figure 4). This resulted in 17 and 5 proteins with statistically significant changes in the UTEX 2973_pEG001 (Table 1) and PCC 6803_pEG001 (Table 2), respectively. Strikingly, the formation of nanofilaments did not lead to many changes in either strain. This is particularly true for PCC 6803 where only 5 proteins were found to have been significantly changed. These findings agree with the observed minor impact on growth (Figure 1B).

**Figure 4.**
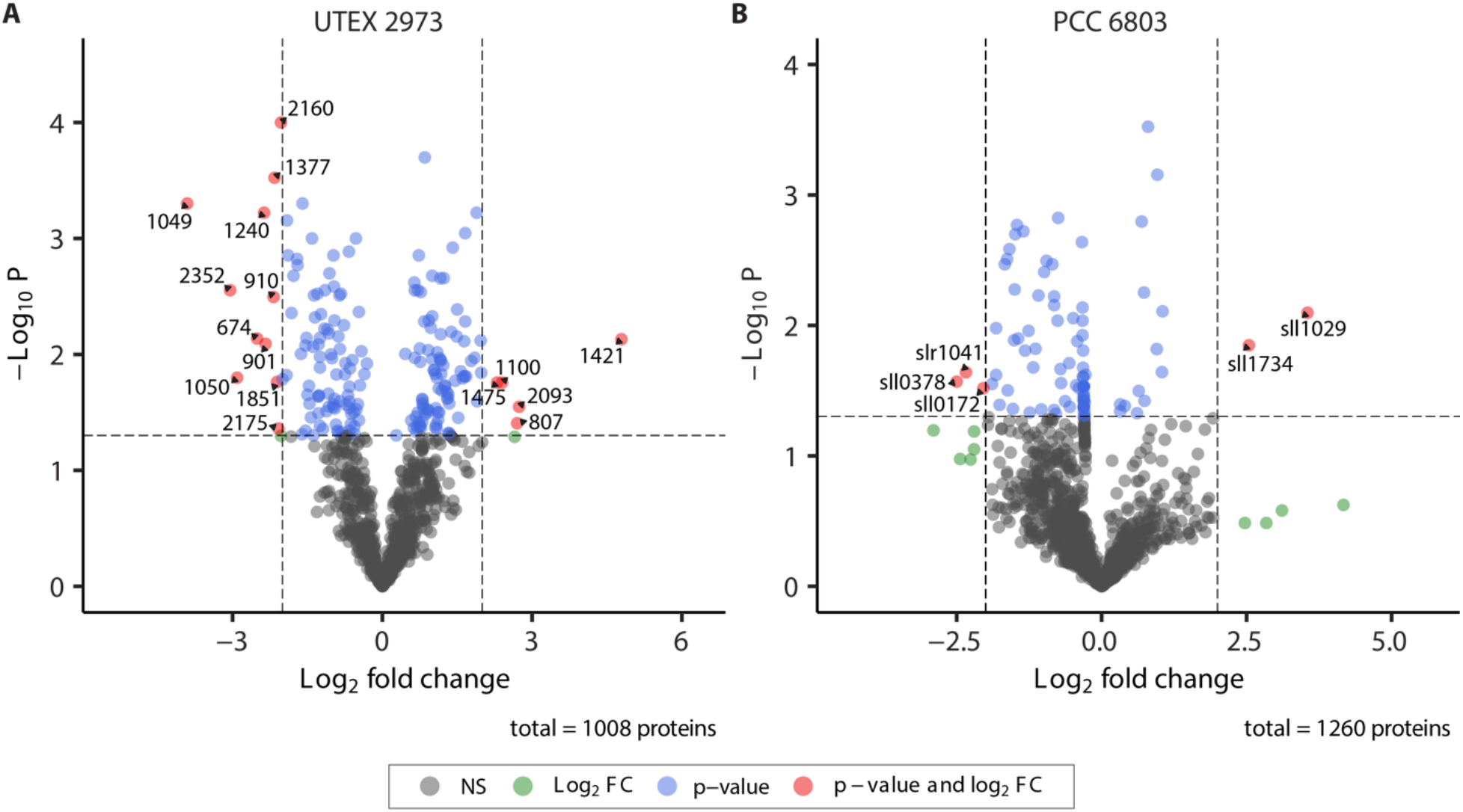
Volcano plots representing proteomic changes in nanofilament forming cyanobacteria. **A, B** The averages of the identified proteins in UTEX 2973_pEG001 (1008 proteins, n = 3) and PCC 6803_pEG001 (1260 proteins, n = 2) were compared to their respective empty vector controls. Red circles show proteins which are significantly up- or downregulated (fold change ≤ −2 and ≥ 2, and p-value < 0.05). Green circles show proteins which fulfill the log change criterium but are not significantly different. Blue circles show proteins which are significantly different but do not fulfill the log change criterium. Black circles show proteins without any differences.

**Table 1.** Differentially expressed proteins in *S. elongatus* UTEX 2973_pEG001 nanofilament forming strain compared to an empty vector control (UTEX 2973_pDF).

| UniProtKB | Locus tag | Protein | FC |
| --- | --- | --- | --- |
| Q03511 | synpcc7942_1421 | CO<sub>2</sub>-concentrating mechanism protein CcmK2 | 4.79 |
| Q8VPV7 | synpcc7942_2093 | CO<sub>2</sub> hydration protein | 2.74 |
| Q31Q30 | synpcc7942_0807 | DUF4351 domain-containing protein | 2.70 |
| Q31P89 | synpcc7942_1100 | Chromosomal replication initiator protein DnaA | 2.40 |
| Q31N64 | synpcc7942_1475 | Sodium-dependent bicarbonate transporter | 2.30 |
| Q31PD8 | synpcc7942_1051 | Phycobilisome rod linker polypeptide | -2.00 |
| Q31L79 | synpcc7942_2160 | Alanine-glyoxylate aminotransferase | -2.03 |
| Q31L64 | synpcc7942_2175 | Iron deficiency-induced protein A | -2.08 |
| Q31M38 | synpcc7942_1851 | Ferredoxin–nitrite reductase | -2.11 |
| Q31NG2 | synpcc7942_1377 | Metal dependent phosphohydrolase | -2.16 |
| Q31PS9 | synpcc7942_0910 | Uncharacterized protein | -2.18 |
| Q31PT8 | synpcc7942_0901 | Haloalkane dehalogenase | -2.34 |
| P39661 | synpcc7942_1240 | Ferredoxin–nitrite reductase | -2.37 |
| Q31QG3 | synpcc7942_0674 | Oxygen-dependent coproporphyrinogen-III oxidase | -2.51 |
| Q31PD9 | synpcc7942_1050 | Phycobilisome rod linker polypeptide | -2.91 |
| Q31KN7 | synpcc7942_2352 | Ribosome hibernation promoting factor (HPF) | -3.05 |
| Q31PE0 | synpcc7942_1049 | Phycobilisome rod linker polypeptide | -3.91 |

**Table 2.** Differentially expressed proteins in *Synechocystis* sp. PCC 6803_pEG001 nanofilament forming strain compared to an empty vector control (PCC 6803_pDF).

| UniProtKB | Locus tag | Protein | FC |
| --- | --- | --- | --- |
| P72760 | sll1029 | CO<sub>2</sub>-concentrating mechanism protein CcmK1 | 3.55 |
| P73393 | sll1734 | CO<sub>2</sub> hydration protein | 2.54 |
| Q55558 | sll0172 | Unknown protein | -<br>2.04 |
| P73005 | slr1041 | PatA subfamily | -<br>2.34 |
| Q55749 | sll0378 | Uroporphyrinogen-III C-methyltransferase | -<br>2.50 |

In both nanofilament-forming strains, proteins involved in carbon fixation were found to be the most upregulated (Tables 1 and 2). These include carboxysome shell proteins CcmK2 (Synpcc7942_1421) and CcmK1 (Sll1029) and CO_2_ hydration proteins (Synpcc7942_2093 and Sll1734). In UTEX 2973_pEG001, the proteins that were considered downregulated indicate clear signs of nitrogen stress. This includes the strong downregulation of phycobiliproteins, proteins involved in nitrogen metabolism (Synpcc7942_1851), amino acid metabolism (Synpcc7942_2160) and cofactor biosynthesis (Synpcc7942_0674). In PCC 6803_pEG001, the signs of nitrogen stress were not as pronounced with only a single protein involved in cofactor biosynthesis (Sll0378) significantly downregulated. In cyanobacteria, carbon and nitrogen metabolism are sinks for ATP and photosynthetic reducing power and balancing the two is crucial to cellular homeostasis^29^. Therefore, our data suggest that the production and maintenance of the nanofilaments may be causing a C/N imbalance which the cells are trying to overcome. Similar responses have been observed in proteomic studies where cyanobacteria are challenged by different environmental perturbations^30^. In addition, this hypothesis is supported by the fact that, in PCC 6803_pEG001, the formation of smaller nanofilaments led to a decreased metabolic burden. Ultimately, our data suggest that the changes observed can mostly be attributed to the metabolic burden of producing, and maintaining, the nanofilaments.

### Recruitment of heterologous proteins to nanofilaments

Next, we sought to target the fluorescent protein mCitrine to the observed PduA* nanofilaments as a proof-of-concept for the recruitment of cargo proteins. To achieve this, we constructed a replicative plasmid for the co-expression of PduA* and mCitrine, the latter fused to the encapsulation peptide P18. P18 is an N-terminal peptide sequence of PduP, an internal enzyme of the Pdu metabolosome that interacts with the shell protein PduA^31–33^. We constructed plasmids for the co-expression of PduA*\** and mCitrine with and without the P18 encapsulation peptide. Further controls were included where mCitrine or P18-mCitrine were expressed in the absence of PduA*. For all these combinations of PduA* and cargo protein, the same self-replicative plasmid previously used for PduA* expression was applied. An overview of all generated constructs is given in Figure 5A and Supplementary Table 1. Both strains, UTEX 2973 and PCC 6803, were transformed with all mCitrine constructs (pAS001, pAS002, pAS003 and pAS004) and successful transformation of cyanobacteria was verified by colony PCR (Supplementary Figure 1).

**Figure 5.**
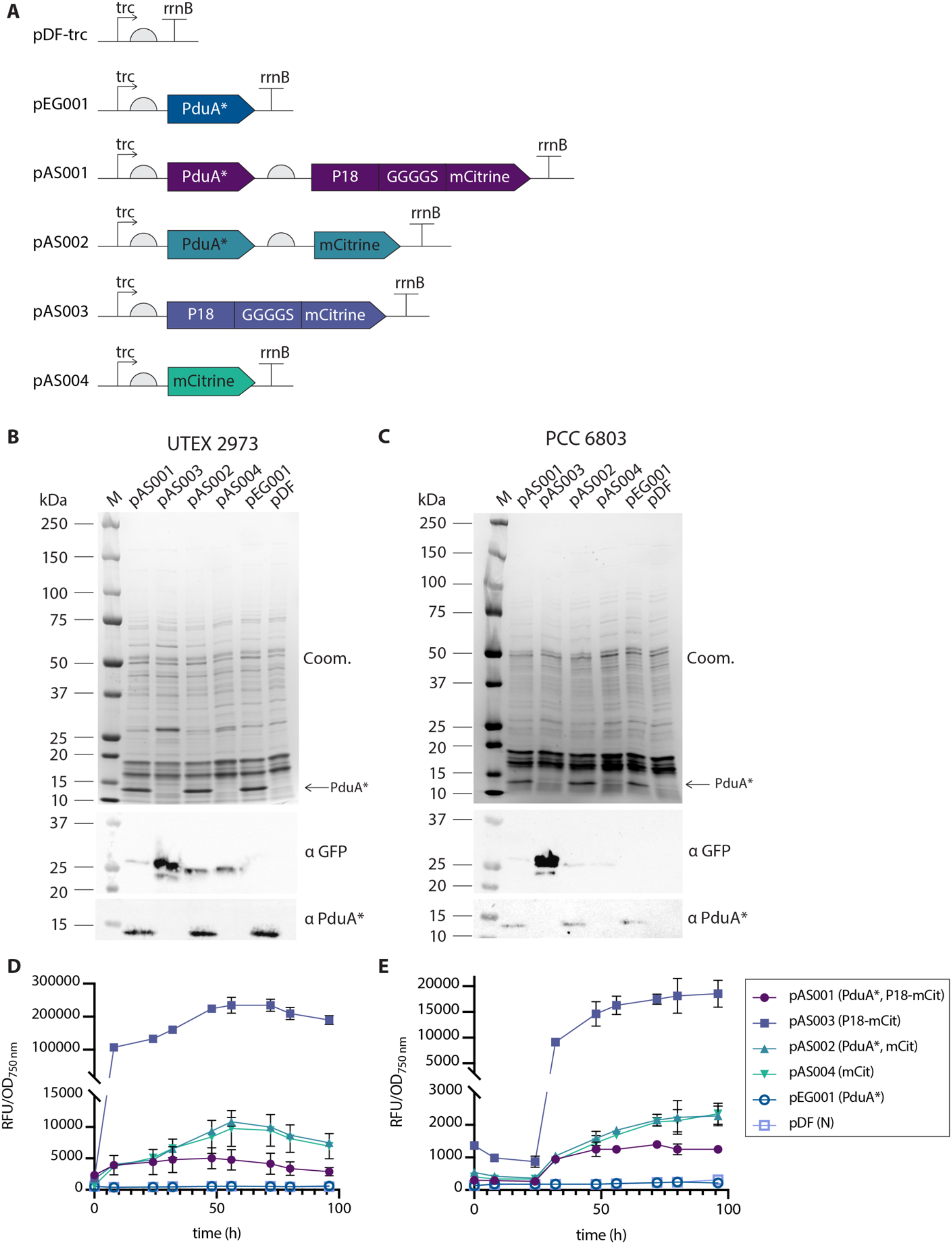
Co-expression of PduA* and mCitrine in *S. elongatus* UTEX 2973 and *Synechocystis* sp. PCC 6803. **A** Overview of construct design in plasmids used in this study. **B, C** Coomassie stain (Coom.) and immunodetection of PduA* (αPduA*) and mCitrine (αGFP) in 5 μg of cleared whole cell lysates of UTEX 2973 (**B**) or PCC 6803 (**C)** strains expressing plasmids p01, pAS002, pAS003, pAS004, pEG001 or pDF-trc. Samples were harvested 48 hours after induction. The PduA* protein band is marked with an arrow. M denotes a molecular weight marker**. D** mCitrine fluorescence (RFU, excitation: 490 nm, emission: 531 nm) in UTEX 2973 strains over a period of 4 days of cultivation and normalized to OD_750 nm_. Data points are averages, error bars represent the standard deviation (n = 3). **E** mCitrine fluorescence (RFU, excitation: 490 nm, emission: 531 nm) in PCC 6803 strains over a period of 4 days of cultivation and normalized to OD_750 nm_. Data points are averages, error bars represent the standard deviation (n = 3).

Expression levels of mCitrine and PduA* in the generated UTEX 2973 and PCC 6803 transformants were analyzed in cleared cell lysates by western blotting (Figure 5B, 5C) and similar trends were found for both groups. All strains containing PduA* (pEG001, pAS001, pAS002) showed similar levels of PduA* expression, however the expression levels of mCitrine varied significantly between strains. High expression levels were detected for P18-mCitrine in the absence of PduA* (UTEX2973_pAS003, PCC 6803_pAS003) while all other combinations of mCitrine showed similar, lower levels of mCitrine expression (Figure 5B, 5C). The expression patterns of mCitrine fluorescence detected by western blotting also matched with the mCitrine fluorescence levels measured in whole cell cultures (Figure 5D, 5E).

To analyze expression of PduA* and mCitrine further, cell growth was recorded over a period of 96 hours (Supplementary Figure 5) and mCitrine fluorescence was simultaneously monitored (Figure 5D, 5E). Strains co-expressing PduA* and P18-mCitrine showed constant levels of fluorescence upon induction of expression in UTEX 2973 (Figure 5D, pAS001) and PCC 6803 (Figure 5E, pAS001). The fluorescence levels of these transformants were consistently lower than in strains expressing mCitrine with and without the encapsulation peptide P18. On the other hand, significantly higher fluorescence was recorded in strains expressing P18-mCitrine in the absence of PduA* (Figure 5, pAS003). This finding is consistent with what was reported when the first nineteen amino acids (P19 instead of P18) of the same encapsulation peptide were fused with a β-galactosidase and an esterase^34^. As previously reported, this strong fluorescence is likely caused by self-aggregation of the encapsulation peptide^12,35^. However, P18-mediated self-aggregation of mCitrine did not occur in the presence of PduA* (Figure 5, pAS001). We hypothesize that PduA* may have served as a competitive binding partner for P18-mCitrine, which supports a P18-mediated interaction of the cargo protein mCitrine and the PduA* nanofilaments.

To investigate the recruitment of P18-mCitrine to PduA* further, we used ultracentrifugation to separate whole cell lysates of cultures co-expressing PduA* and mCitrine with and without P18 (pAS001 and pAS002, respectively) on sucrose gradients from 0 to 60% sucrose. If the P18 peptide facilitates a close interaction of mCitrine with PduA*, we reasoned this would lead to larger protein complexes which partition differently within the sucrose gradient. First, oriole staining of separated sucrose fractions was used to confirm the suitability of the developed protocol (Supplementary Figure 6). Following this, the sucrose fractions were analyzed by immunoblotting. PduA* was detected at sucrose concentrations between 15 and 60% in both strains of UTEX 2973 (Figure 6A) and PCC 6803 (Figure 6B). Similar PduA* distribution patterns were observed in co-expression with both P18-mCitrine (pAS001) and mCitrine (pAS002). In contrast, distinct distribution patterns of mCitrine were observed in the presence or absence of the P18 encapsulation peptide (Figure 6A, B). In UTEX 2973, the addition of P18 led to the migration of mCitrine into higher density fractions. Strikingly, the majority of P18-mCitrine was detected in the 60% sucrose layer where PduA* was also found (Figure 6A). Without the P18 encapsulation peptide, mCitrine was mainly found in the 10% sucrose layer. In PCC 6803, the differences in distribution patterns of mCitrine with and without P18 across the sucrose gradient were less pronounced. Still, the same trend as for UTEX 2973, with P18 leading to partitioning of mCitrine into higher sucrose densities of up to 40% sucrose, was observed (Figure 6B). These findings suggest, in particular for UTEX 2973, that the P18 encapsulation peptide mediates an interaction between mCitrine and PduA* resulting in the formation of larger complexes that migrate into higher sucrose concentrations.

**Figure 6.**
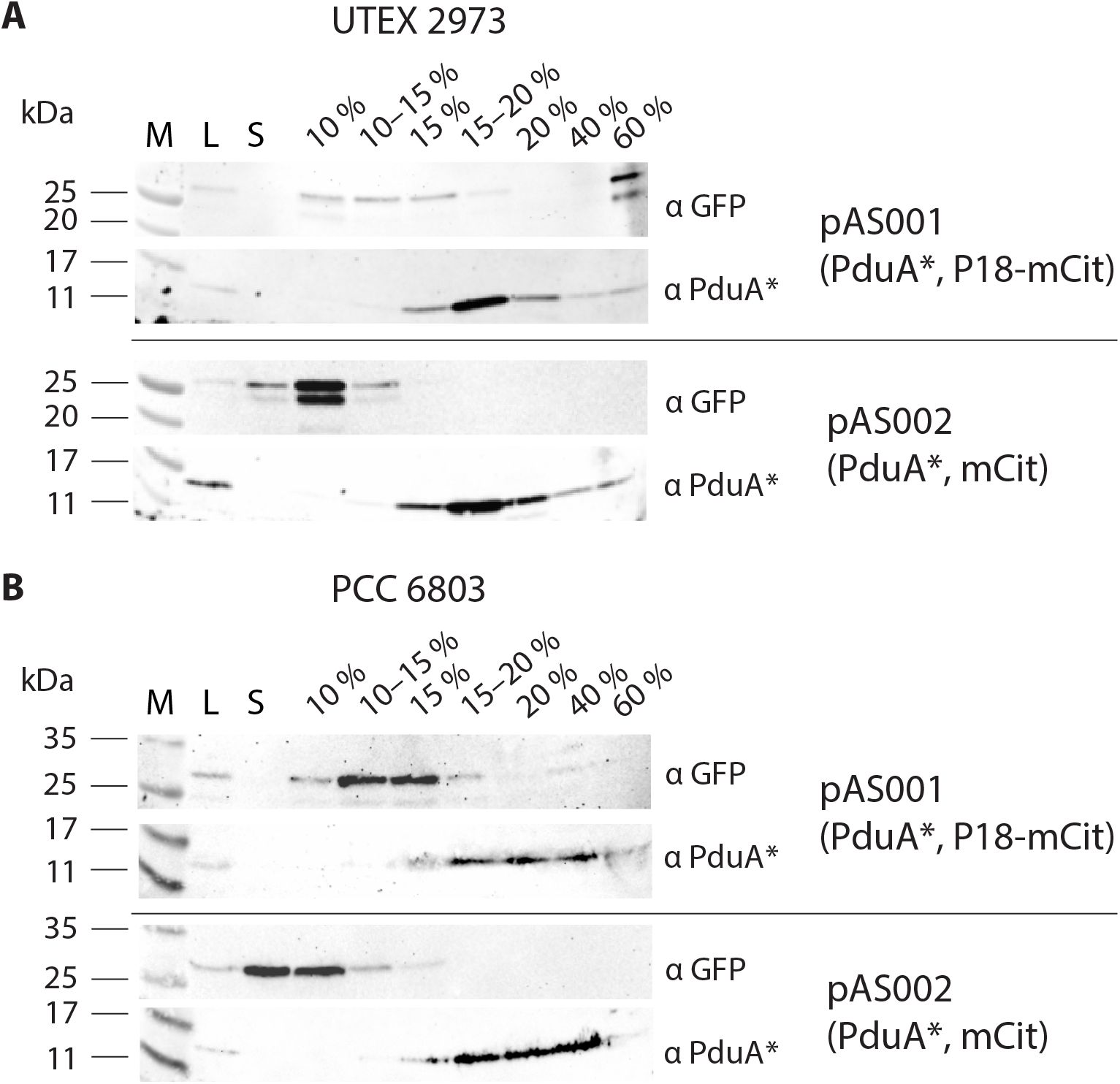
Detection of PduA* and mCitrine in cellular fractions separated by sucrose gradient ultracentrifugation. **A** Immunoblots detecting mCitrine (αGFP) and PduA* (αPduA*) in sucrose gradient separated cellular fractions of UTEX 2973_pAS001 and UTEX2973_pAS002. **B** Immunoblots detecting mCitrine (αGFP) and PduA* (αPduA*) in sucrose gradient separated cellular fractions of PCC 6803_pAS001 and PCC6803_pAS002. M: molecular weight marker, L: 2.5 μg of cleared whole cell lysate before fractionation, S: sucrose gradient sample layer. Percentages of lanes refer to the sucrose density of the individual sucrose layer.

In summary, the P18 encapsulation peptide was found to enable interaction of the fluorescent reporter mCitrine with the PduA* nanofilaments. However, several shortcomings of this delivery system will need to be addressed in future work. The interaction of a cargo protein with the PduA* filaments mediated by encapsulation peptides was found to be rather weak in *E. coli* and the self-aggregating properties of P18 compete with cargo-nanofilament interactions^35^. Our data suggest a similar scenario in cyanobacteria. Several routes of optimization to improve the cargo-nanofilament protein interactions could be explored. For instance, this could be achieved using synthetic coiled-coil peptides^12^ or by repurposing native cyanobacterial proteins rich in coiled-coils^36^. Furthermore, a recent study reports that approximately 20% of bacterial genomes encode BMCs with a wide phylogenetic distribution^37^. Tapping into this diversity of native BMC technology, in conjunction with advances in understanding BMC technology, has great potential to optimize heterologous expression systems including cyanobacterial cell factories^38^.

## Conclusions

In this study we successfully engineered PduA*-based nanofilaments into two different cyanobacteria: the model organism PCC 6803 and the fast-growing strain UTEX 2973. PduA* was found to form structures resembling nanofilaments in UTEX 2973 and nanofilaments or sheets in PCC 6803. Shotgun proteomics of the nanofilament-forming strains suggested only a minor metabolic impact on cellular metabolism, mainly associated with nutrient stress. Finally, we demonstrated that a cargo protein can be recruited to the filaments using the fluorescent protein mCitrine in conjunction with the encapsulation peptide P18. This proof-of-concept study takes a significant step towards unlocking the potential of nanobiotechnology in photosynthetic production systems. In addition, it paves the way for the development of future applications such as the organization of heterologous pathways in cyanobacteria. Ultimately, protein-based nanofilaments hold great potential as a tool to increase the production efficiency and yields of cyanobacterial cell factories.

## Author Contributions

Concept and Experimental Design: JAZZ, SF, PEJ. Experimental Work and Data Collection: Construct Design – JAZZ; Strain generation – AMS, EG; Growth and fluorescence analysis –AMS; Immunoblotting –JAZZ, AMS, EG; Sucrose fractionation: AMS; Proteomics: JAZZ, DAR, EG, AM; transmission electron microscopy: JAZZ, LH, PV. Data analysis: JAZZ, AMS, DAR, EG, AM, SF, PV, PEJ. Manuscript Writing: JAZZ, AMS, DAR. All authors have read and approved the final version of the manuscript.

## Acknowledgments

The authors thank Dr. Chris Neal and Judith Mantell of the Wolfson Bioimaging Facility of the University of Bristol for their expert assistance with transmission electron microscopy, Prof. Mike Jones and Dr. Paul Curnow for providing JAZZ access to their facilities at the University of Bristol, Prof. Martin Warren and Dr. Matthew Lee for providing the PduA* DNA sequence and Dr. Christoph Crocoll for help with data management. The authors acknowledge financial support from the ‘Cynthetica’ project from the European Union’s Horizon 2020 research and innovation programme under the Marie Skłodowska-Curie grant agreement no. 745959 (JAZZ), the Humboldt Foundation (DAR), BBSRC sLoLa Research grant (BB/M002969/1) (PV), the Novo Nordisk Foundation (NNF19OC0057634) (PEJ) and the Carlsberg Foundation (CF17-0657) (PEJ).

## Supplementary Material

**Supplementary Information:** colony PCR of generated transformants; electron micrographs; growth curves; oriole stain of cellular lysates separated by sucrose gradient ultracentrifugation; lists of plasmids and primers used in this study.

**Supplementary Data 1:** list of proteins identified in UTEX 2973 and PCC 6803 expressing pEG001 and empty vector controls.

**Supplementary Video 1, 2, 3:** Electron tomograms of UTEX2973_pEG001 strain.

