## Supplementary Information for "Self-assembly of nanofilaments in cyanobacteria for protein co-localization"

### Supplementary Information
### Self-assembly of nanofilaments in cyanobacteria for protein co-localization

*Julie A. Z. Zedler[1‡*], Alexandra M. Schirmacher[1‡], David A. Russo[2], Lorna Hodgson[3], Emil Gundersen[4], Annemarie Matthes[4], Stefanie Frank[5], Paul Verkade[3] and Poul Erik Jensen[6]*

[1]Synthetic Biology of Photosynthetic Organisms, Matthias Schleiden Institute for Genetics, Bioinformatics and Molecular Botany, Friedrich Schiller University Jena, Jena, Germany
[2]Bioorganic Analytics, Institute for Inorganic and Analytical Chemistry, Friedrich Schiller University Jena, Jena, Germany
[3]School of Biochemistry, University of Bristol, Bristol, United Kingdom
[4]Department of Plant and Environmental Sciences, University of Copenhagen, Frederiksberg, Denmark
[5]Department of Biochemical Engineering, University College London, London, United Kingdom
[6]Department of Food Science, University of Copenhagen, Frederiksberg, Denmark.

‡these authors contributed equally.

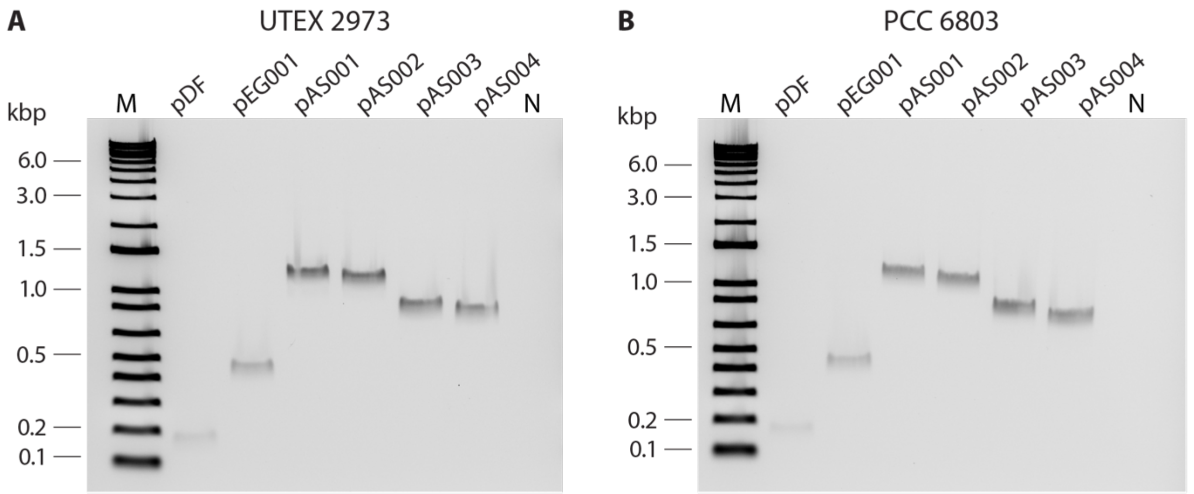


**Supplementary Figure 1.** Colony PCR of generated cyanobacterial strains using the primer pair “trc-F” and “trc-R” as previously described by Russo et al.1 **A** *Synechococcus elongatus* UTEX 2973 and **B** *Synechocystis* sp. PCC 6803 transformed with plasmids pDF, pEG001, pAS002, pAS003 and pAS004, respectively. M: molecular weight marker in kilo base pairs. N: PCR negative control reaction using water.*, further confirmed that this highly abundant protein was PduA* (Figure 1A).

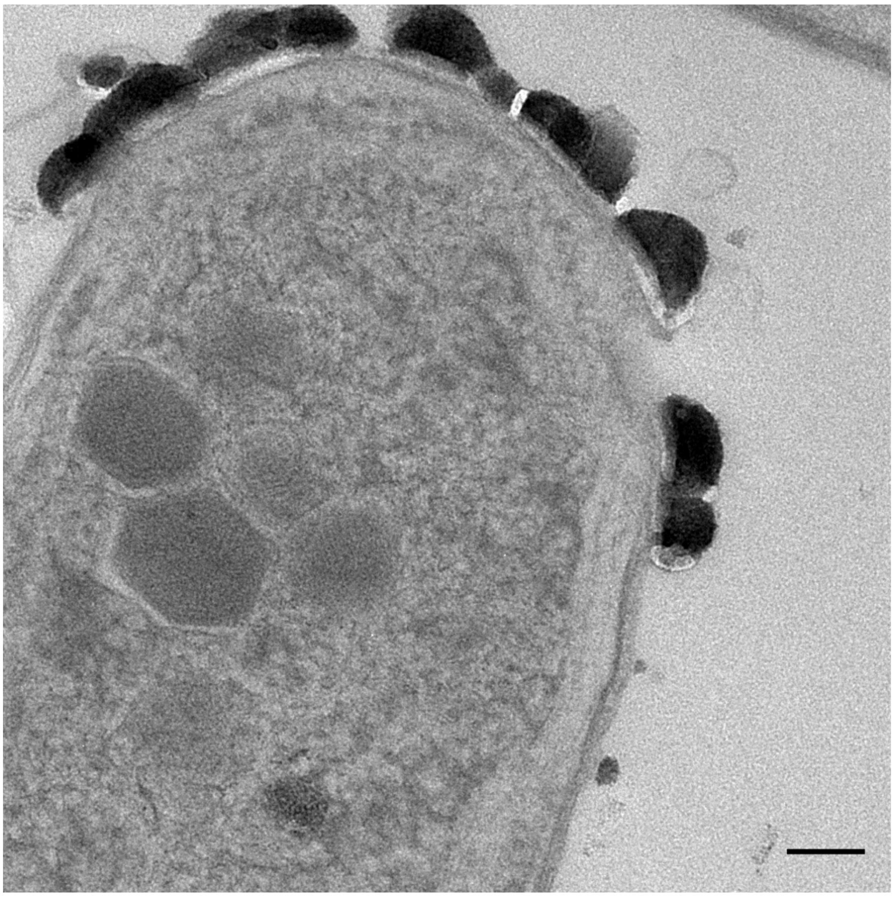

**Supplementary Figure 2.** Electron micrograph of *Synechococcus elongatus* strain UTEX 2973_pDF. Scale bar: 100 nm.

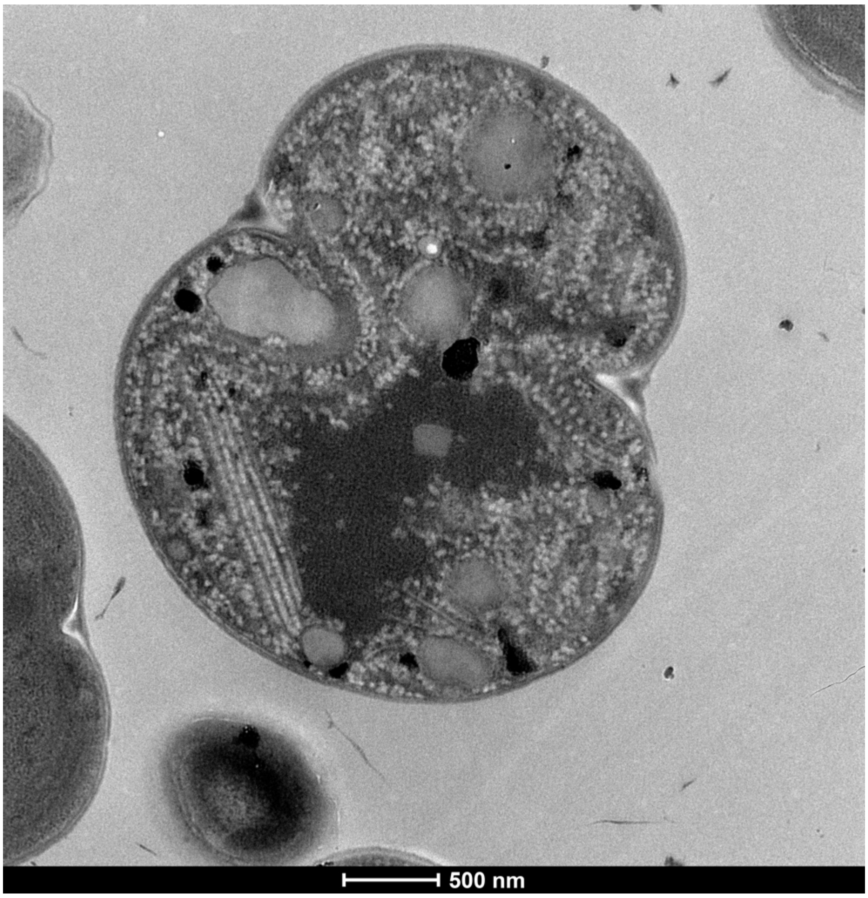


**Supplementary Figure 3.** Electron micrograph of *Synechocystis* strain PCC 6803_pEG001. Scale bar: 500 nm.

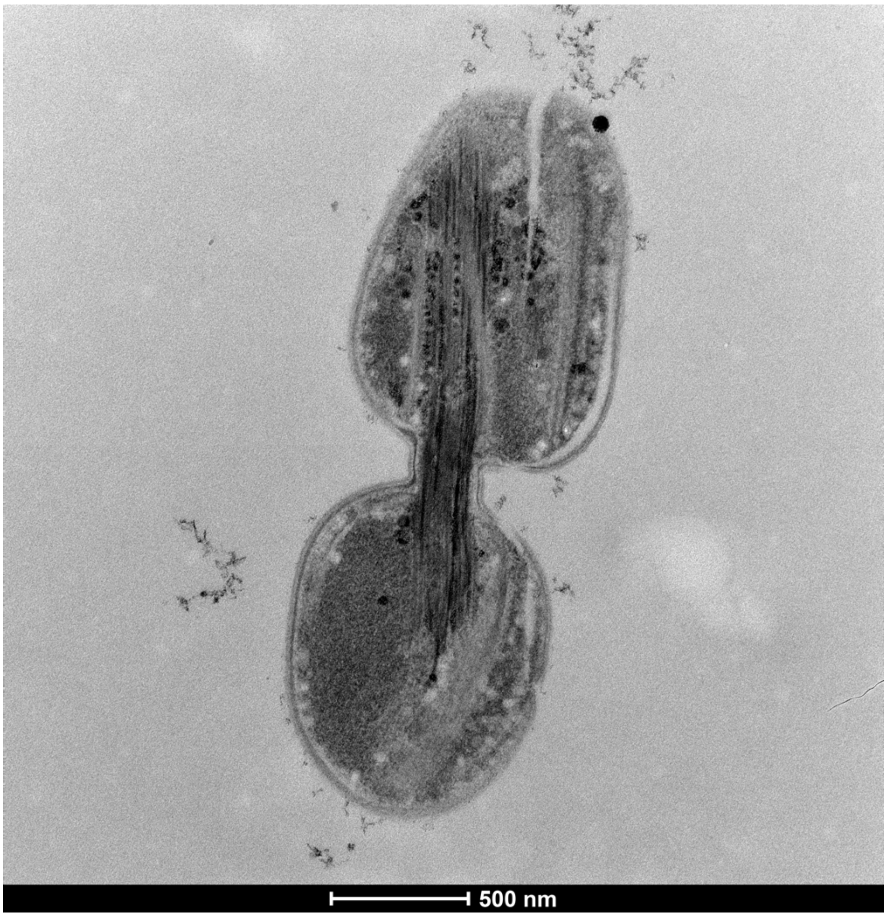


**Supplementary Figure 4.** Electron micrograph of *Synechococcus elongatus* strain UTEX 2973_pEG001. Scale bar: 500 nm.

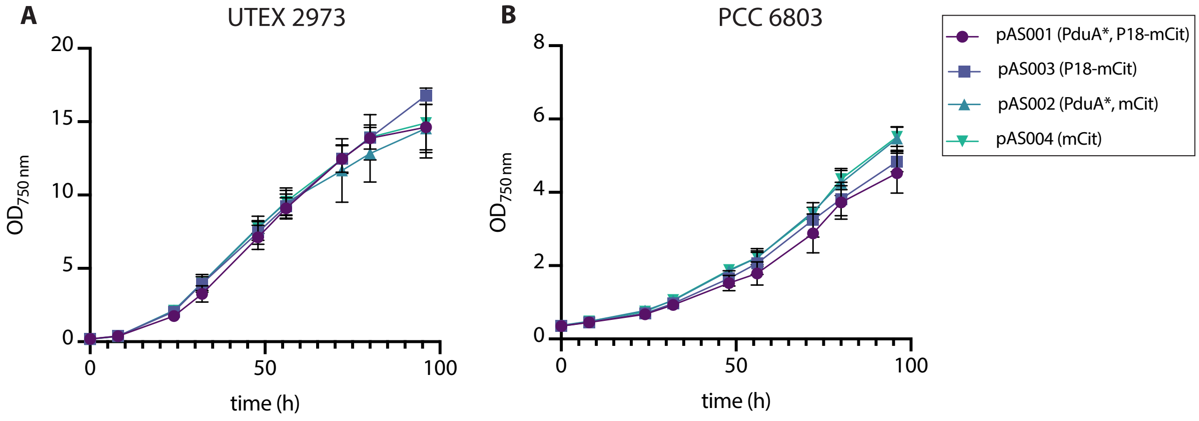


**Supplementary Figure 5.** Growth of strains expressing PduA* and (P18-)mCitrine over a period of 4 d. **A** $OD_{750nm}$ of UTEX 2973 strains expressing plasmids pAS001, pAS002, pAS003 and pAS004 grown at 30°C, 720 µmol photons $m^{-2}$ $s^{-1}$ with 3% $CO_2$-enriched air bubbling. **B** $OD_{750nm}$ of PCC 6803 strains expressing plasmids pAS001, pAS002, pAS003 and pAS004 grown at 30°C, 75 µmol photons $m^{-2}$ $s^{-1}$ with 3% $CO_2$-enriched air bubbling.

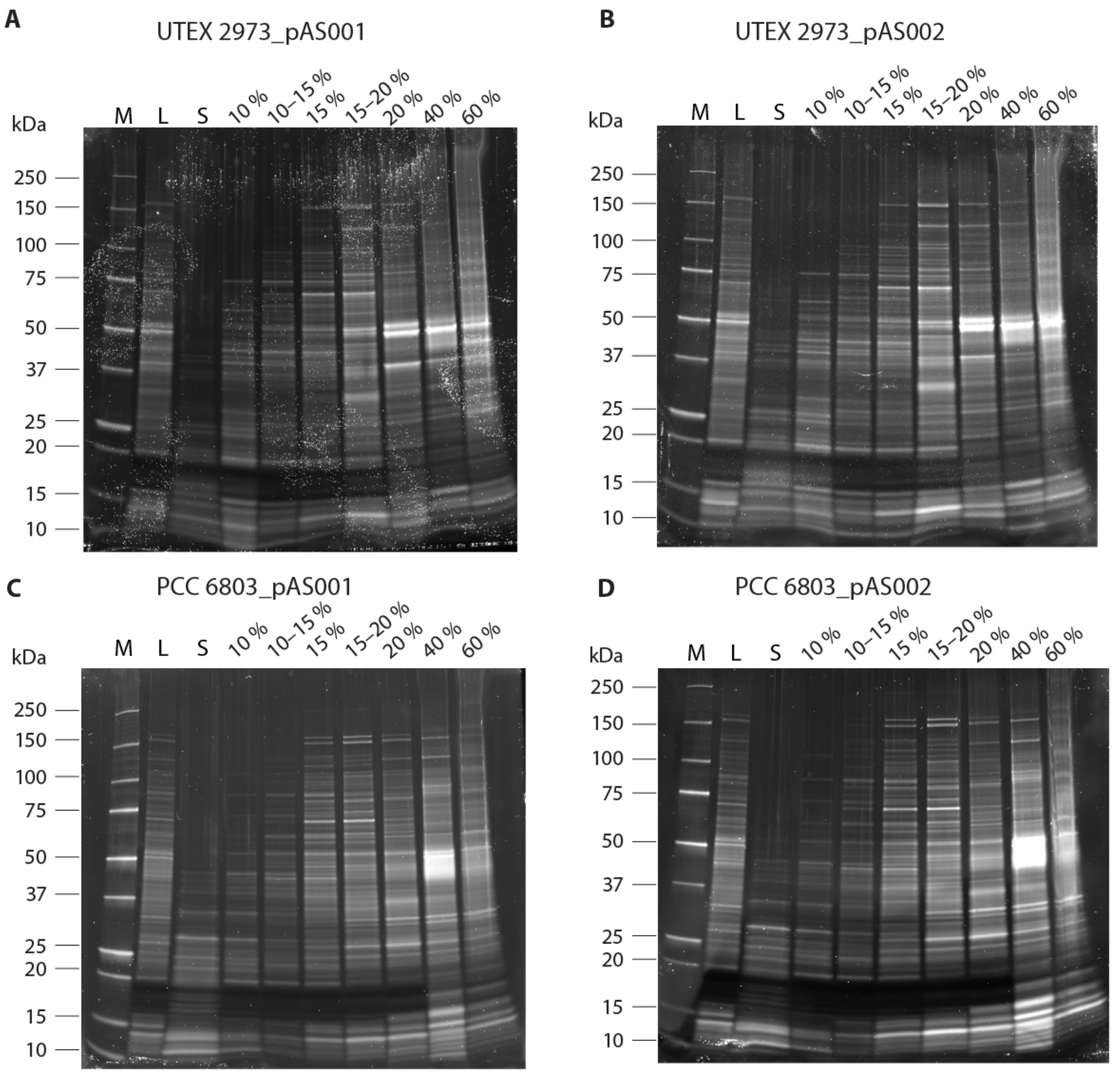


**Supplementary Figure 6.** Oriole stain of cellular lysates separated by sucrose gradient ultracentrifugation. **A** Fractions from UTEX 2973_pAS001 (PduA*, P18-mCitrine), **B** UTEX 2973_pAS002 (PduA*, mCitrine), **C** PCC 6803_pAS001 (PduA*, P18-mCitrine) and **D** PCC 6803_pAS002 (PduA*, mCitrine). M: molecular weight marker, L: whole cell lysate loaded onto sucrose gradients, S: sucrose gradient sample layer. Percentages of lanes refer to the sucrose density of the individual sucrose layer or interphase respectively.

**Supplementary Table 1.** Plasmids used in this study.

| Plasmids | Details | Origin of this plasmid |
|---|---|---|
| pDF-trc | Replicative plasmid; selection marker: Streptomycin resistance cassette. | Guerrero et al., 2012[2] |
| pEG001 | Ptrc, pduA*, TrrnB; backbone: pDF-trc. | this study |
| pAS001 | Ptrc, pduA*, p18-mCitrine, TrrnB; backbone: pEG001. | this study |
| pAS002 | Ptrc, pduA*, mCitrine, TrrnB; backbone: pEG001. | this study |
| pAS003 | Ptrc, p18-mCitrine, TrrnB; backbone: pDF-trc. | this study |
| pAS004 | Ptrc, mCitrine, TrrnB; backbone: pDF-trc. | this study |

**Supplementary Table 2.** Primers used for construct assembly.

| Primer | Sequence (5'-3') | Used for generating the following plasmids |
|---|---|---|
| PduA.F1 | CGTCGAGAATTCATGCAACAAGAAGCGTTAGGAATG | pEG001 |
| PduA.R1 | GTAGACAAGCTTTTATTGCTCAGCGGTGGC | pEG001 |
| G.pDF_mCit R | TCCGCCAAAACAGCCAAGCTTCACTTGTACAGCTCGTCCA | pAS001 + pAS002 + pAS003 + pAS004 |
| G.pDF_PduA-P18-mCit F | ACCGCTGAGCAATAAAAGCTATTAATAAGCTTTAGTGGAGGTGGATC | pAS001 |
| G.pDF_PduA-mCit F | ACCGCTGAGCAATAAAAGCTATTAATAAGCTTTAGTGGAGGTGGATCCATGGTGAGCAAGGGCGAGG | pAS002 |
| G.pDF_P18-mCit F | AGGAAACAGACCATGGAATTCATGAACACTTCTGAGCTGGAAAC | pAS003 |
| G.pDF_mCit F | AGGAAACAGACCATGGAATTCATGGTGAGCAAGGGCGAGGAG | pAS004 |
